# Ferrous iron uptake via IRT1/ZIP evolved at least twice in green plants

**DOI:** 10.1101/2022.07.21.501042

**Authors:** Wenderson Felipe Costa Rodrigues, Ayrton Breno P. Lisboa, Joni Esrom Lima, Felipe Klein Ricachenevsky, Luiz-Eduardo Del-Bem

## Abstract

Iron (Fe) is an essential micronutrient for virtually all living beings, being practically irreplaceable because of its unique electrochemical properties that enable or facilitate a series of biochemical processes, including photosynthesis. Although Fe is abundant on Earth, it is generally found in the poorly soluble form Fe^3+^. Most extant plants have established Fe absorption strategies that involve Fe uptake in the soluble form Fe^2+^. The model angiosperm *Arabidopsis thaliana*, for example, captures Fe through a mechanism that lowers the pH through proton pumping to the rhizosphere to increase Fe^3+^ solubility, which is then reduced by a plasma membrane-bound reductase and transported into the cell by the ZIP family protein IRT1. ZIP proteins are transmembrane transporters of a variety of divalent metals such as Fe^2+^, Zn^2+^, Mn^2+^ and Cd^2+^. In this work, we investigate the evolution of functional homologs of IRT1/ZIP in the supergroup of photosynthetic eukaryotes Archaeplastida (Viridiplantae + Rhodophyta + Glaucophyta) using a dataset of 41 high-quality genomes of diverse lineages. Our analyses suggest that Fe is acquired through deeply divergent ZIP proteins in land plants and chlorophyte green algae, indicating that Fe^2+^ uptake by ZIP family proteins evolved at least twice independently during green plant evolution. Sequence and structural analyses indicate that the archetypical IRT proteins from angiosperms likely emerged in streptophyte algae before the origin of land plants and might be an important player in green plant terrestrialization, a process that involved the evolution of Fe acquisition in terrestrial subaerial settings.

## 1. Introduction

Iron (Fe) is a fundamental element for virtually all cellular organisms. In plants, Fe deficiency results in serious physiological problems, such as failures in photosynthetic electron transport and sulfur and nitrogen metabolism, resulting in impaired growth (Crichton, 2016). Fe is not readily bioavailable, besides being one of the most abundant metals in Earth’s crust. In soil, Fe is predominantly found as Fe^3+^ (ferric iron), which is poorly bioavailable to plants. In opposition, Fe^2+^ (ferrous iron) can be readily absorbed by plant roots without being reduced and without the aid of phytosiderophores. In freshwater environments, Fe can be found dissolved in greater abundance than in marine environments, where its limitation impacts photosynthetic productivity (Frey and Reed, 2012; Blomqvist et al., 2004).

The great physiological importance that Fe has for plants and other living organisms is possibly explained by the role of Fe-based catalysts in early life, before the origin of extant cellular domains (Camprubi et al., 2017). The decrease of Fe bioavailability during the Great Oxidation Event, caused by the emergence of oxygenic photosynthesis, likely shaped the evolution of life afterward (Wade et al., 2021). Throughout evolutionary time, organisms acquired specific mechanisms for Fe uptake that can sense and respond to soluble Fe in the environment. Land plants (Embryophyta) evolved at least two different Fe capture strategies. Based on Fe reduction, Strategy I is carried out by most plants, and Strategy II, based on chelation, seems to be specific to the Poaceae family (Römheld and Marschner, 1986; Connorton et al., 2017).

In *Arabidopsis thaliana* Strategy I (Figure 1a), root cells pump protons to the surrounding soil through H-ATPase 2 (AHA2), decreasing the pH of the rhizosphere and increasing the Ferric Fe hydroxide (Fe(OH)_3_) solubility. Fe^3+^ is then reduced by FRO2 (Ferric Reduction Oxidase 2) reductase to Fe^2+^ and transported into cells by the ZIP (Zinc-regulated, Iron-regulated transporter-like Protein) transporter called IRT1 (Iron-Regulated Transporter 1) (Robinson et al., 1999; Henriques et al., 2002; Varotto et al., 2002; Vert et al., 2002; Santi and Schmidt, 2009). Low availability of Fe induces the synthesis of AHA2, FRO2, and IRT1 proteins that form complexes at the plasma membrane of the root epidermis cells (Martín-Barranco et al., 2020). The acidification-reduction step can also be performed using Fe-mobilizing coumarins, such as fraxetin, sideretin and esculetin (Kai et al., 2006; Rajniak et al., 2018; Siwinska et al., 2018; Tsai et al., 2018; Robe et al., 2021). The F6’H1 (Feruloyl-CoA 6’-Hydroxylase 1) enzyme is responsible for the conversion of feruloyl CoA to 6’-hydroxyferuloyl CoA which is the common precursor for the synthesis of coumarins secreted in the rhizosphere by ABCG transporter PDR9 (Pleiotropic Drug Resistance 9) (Rodriguez-Celma et al., 2013; Fourcroy et al., 2014; Schmid et al., 2014). Prior to their secretion into the rhizosphere, coumarins are deglycosylated by β-glucosidases (BGLUs) (Clemens and Weber, 2016; Connorton et al., 2017).

**Figure 1.**
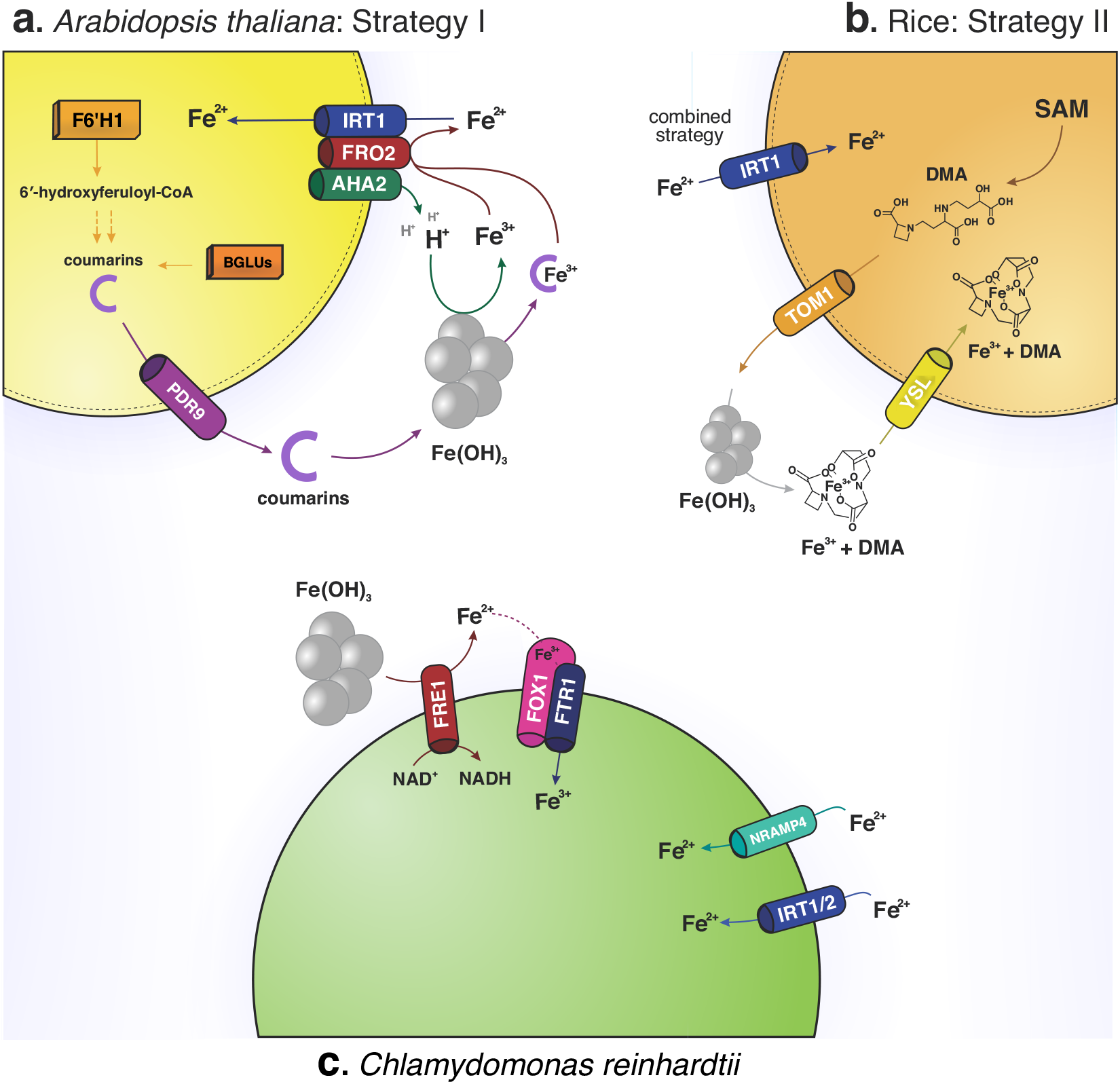
Iron uptake strategies in (a) Arabidopsis thaliana (Strategy I), (b) rice (Strategy II and IRT1-dependent Fe2+ uptake), and (c) Chlamydomonas reinhardtii. Cylindrical shapes indicate membrane proteins involved in different steps of Fe capture. Blue shapes represents Fe transporters and red shapes represent reductases. Orange boxes inside cells represents enzymes involved in the biosynthesis of coumarins.

In Strategy II, root cells secrete phytosiderophores into the soil, which chelate Fe(OH)3 (Takagi et al., 1984). Phytosiderophores are synthesized from S-adenosyl-methionine (SAM), generating compounds of the mugineic acid (MA) family, such as 2’-deoxymugineic acid (DMA), that are secreted via TOM1 (Transporter of Mugineic acid family phytosiderophores 1) (Takagi et al., 1984; Mori and Nishizawa, 1987; Cheng et al., 2007; Nozoye et al., 2011; Bashir et al., 2017). Phytosiderophores form complexes with Fe^3+^ that are transported into the cell by YSL (Yellow Stripe1-Like) (Figure 1b) (Inoue et al., 2009; Lee et al., 2009). Species of the rice genus *(Oryza)* can combine Strategy II with direct uptake of Fe^2+^ via IRT1, in what is called the Combined Strategy. This is probably an adaptation of roots to flooded areas, where the lower O2 levels causes Fe to be reduced to Fe ^2+^ (Connorton et al., 2017; Wairich et al., 2019; Martín-Barranco et al., 2021).

In green algae, such as *Chlamydomonas reinhardtii* (Chlorophyta), Fe absorption occurs by reduction through FRE1 reductase (Ferric/cupric reductase transmembrane component 1), followed by reoxidation by the putative multicopper oxidase FOX1. Fe is then transported into the cell by the FTR1 permease (high affinity iron permease 1). The FOX1–FTR1 complex is a high-affinity Fe acquisition system that acquires Fe under very low concentrations. *Chlamydomonas* also directly captures Fe^2+^ available in freshwater through the ZIP transporters CrIRT1 and CrIRT2, and the metal ion transporter CrNRAMP4 (Natural Resistance-Associated Macrophage Protein 4) (Figure 1c) (Allen et al., 2007; Blaby-Haas and Merchant, 2012; Glaesener et al., 2013; Kroh and Pilon, 2020; Martín-Barranco et al., 2021).

IRT1 is the first described and most well-known member of the large ZIP family of membrane transporters, identified from an *Arabidopsis* cDNA capable of complementing the Fe uptake function in yeast mutants (Eide et al., 1996). ZIP proteins can transport a broad range of metal other than Fe, such as Zinc (Zn) and Manganese (Mn). In fact, in addition to IRT1, the members that gave the ZIP family its name are the yeast Zn transporters AtZRT1 and AtZRT2 (Eide et al., 1996; Milner et al., 2013). There are not many studies on the physiological roles of the other 12 *Arabidopsis ZIP* genes (Milner et al., 2013; Zheng et al., 2018; Pita-Barbosa et al., 2019; Lee et al., 2021), despite their possible importance in the transport of metal ions. IRT1 from rice is encoded by a gene closely related to *IRT1* from *Arabidopsis* suggesting that the IRT1 -based Fe^2+^ uptake mechanism might be ancestral to angiosperms (Ricachenevsky and Sperotto, 2014).

Freshwater and terrestrial/subaerial environments have different Fe distribution and bioavailability. Thus, the evolution of Fe capture from Earth’s crust and early soils may have been one of the crucial steps in the terrestrialization of early unicellular streptophytes that ultimately gave rise to land plants. It is interesting to think about how and when the Fe capture mechanisms of extant green plants evolved to deal with Fe deficiency. Given that an IRT1-based mechanism for Fe^2+^ is present in angiosperms and Chlamydomonas, it was assumed that this mechanism shares a common evolutionary origin in land plants. Based on these questions, this study used a comparative genomics approach to better understand the origin and diversification of IRT1/ZIP proteins in Archaeplastida (Supplemental Figure 1), the supergroup of photosynthetic eukaryotes that comprises the land plants, green and red algae, and the small group of the glaucophytes. Our results suggest that IRT1-based mechanisms of Fe^2+^ capture evolved at least twice in green plants, from genes belonging to remarkably distinct branches of the ZIP family tree. IRT1 homologs responsible for Fe^2+^ transport in the Strategy I of land plants and those involved in the uptake of Fe^2+^ from water in chlorophytes likely evolved independently from deeply divergent ancestral *ZIP* genes.

## 2. Methods

### 2.1 Datasets

The HMM (hidden Markov model) profile for Zip family (PF02535) was obtained from the Pfam database (https://pfam.xfam.org/) (Mistry et al., 2021). The complete predicted proteomes of the analyzed organisms were collected from the databases detailed in Supplemental Table 1.

### 2.2 Identification of ZIP family homologs

We performed a HMMER v3.2.1 (Eddy, 2011) search with default e-value to identify ZIP family proteins in Archaeplastida. We kept only the most representative protein isoform encoded by each unique gene, defined as the most complete isoform. Through an in-house script written in Perl, we discarded partial sequences or pseudogenes by eliminating sequences shorter than 138aa, representing 40% of the length of the well-characterized IRT1 protein of *Arabidopsis thaliana.* Also, we removed 100% redundant sequences with *skipredundant* from EMBOSS v6.6.0 package (Rice et al., 2000).

All subsequent analyses were performed with the filtered sequences and results were plotted in different graphs with RStudio (R Core Team, 2021). The relative genomic frequency of *ZIP* genes per genome was calculated by dividing the number of homologs found in each genome by the total number of protein-coding genes. P-values were calculated for each contrast using the Wilcoxon statistical test in RStudio and p-values > 0.05 were considered not significant.

### 2.3. Sequence alignment

Protein sequences were aligned using MAFFT (https://mafft.cbrc.jp/alignment/software/) (Katoh, 2002) with the following parameters *--thread 10 --reorder --leavegappyregion --maxiterate 1000 --retree 1 --localpair*. The alignment output was exported in Phylip format *(--phylipout*) for subsequent phylogenetic analyses. We also aligned our sequences using MUSCLE in MEGA 11 software (Kumar et al., 2018) to analyze the conservation of amino acids between proteins and calculate the genetic *p*-distance. This analysis was performed using sequence logos generated by WebLogo 3 (http://weblogo.threeplusone.com/) (Crooks et al., 2004), with standard parameters, except for the y-axis unit which was relative frequency instead of bits.

### 2.4 Phylogenetic analysis

The phylogenetic analyses were performed using the maximum-likelihood method implemented in IQTree v1.6.12 (Nguyen et al., 2015). The amino acid substitution model was selected using the SMS method with Akaike Information Criterion (AIC) as the selection criterion. The Best-fit model chosen according to AIC was WAG+R10. The initial tree was calculated with BIONJ and improved by the tree topology search method NNI (Nearest Neighbor Interchanges). The branch support values were calculated with the likelihood-based method aLRT SH-like.

### 2.5 Sequence Similarity Network (SSN)

To corroborate our phylogenetic results, we performed a network analysis based on sequence similarity. We generated these networks with a threshold equal to 40 in the Enzymatic Similarity Tool (EFI-EST tool) (https://efi.igb.illinois.edu/efi-est/) (Zallot et al., 2019). After that, we visualize and edit the network with Cytoscape v3.8.2 software (Shannon et al., 2003), keeping only nodes and edges with identity ≥ 50%.

### 2.6 Protein tridimensional structure modeling and analyses

Signal peptide was predicted before tridimensional structure modeling using Phobius (Kall et al., 2007) and if a signal peptide was predicted, the correspondent sequence was removed prior to modeling (Supplemental Table 2). Protein tridimensional structure modeling was performed using AlphaFold (Jumper et al., 2021), with default parameters. PDB files underwent sidechain energy minimization on FoldIt Standalone (Kleffner et al., 2017), which uses RosettaCM algorithms (Kuhlman et al., 2003), to resolve sidechain atomic clashes on the models. PDB files were then visualized on PyMOL (Schrödinger, 2015).

To assess the root-mean-square deviation (RMSD) of alpha carbon (Cα) atomic positions, we aligned the tridimensional models with RaptorX Structure Alignment Server DeepAlign (Wang et al., 2013) for three or more structures. For pairwise alignment, structures were aligned on PyMOL.

## 3. Results & Discussion

### 3.1 Distribution of IRT1/ZIP family homologs in the plant lineage

In our analyses, we found 492 homolog proteins (Supplemental Table 3), each corresponding to a unique genomic *locus,* distributed across the 41 species of archaeplastidians in our dataset (Supplemental Figure 1). We detected IRT1/ZIP homologs not only in all lineages of green plants but also in the glaucophyte and the rhodophytes, which indicate that genes encoding IRT1/ZIP proteins in land plants were likely inherited vertically from the last common ancestor of Archaeplastida (Figure 2a). Our data suggest that archaeplastidian algae typically have less than 10 *ZIP* genes (Figure 2a). We separated the algae by habitat (marine, freshwater or terrestrial) and noticed a tendency of freshwater algae to display a higher number of *ZIP* genes when compared to terrestrial or oceanic (Supplemental Figure 2a), which could indicate an evolutionary advantage to have an increased repertoire of metal ion transporters. In freshwater environments, algae adopt specific approaches to Fe absorption, such as those described in *Chlamydomonas* (Figure 1c). The solubility of Fe^2+^ is generally temperature-, pH-and oxygen-dependent. In the ocean, when the water is warm and oxygenated, Fe^2+^ is stable for only a few seconds, and can last for hours or days when the temperature and oxygenation are reduced. Much of the ocean is well oxygenated and Fe^2+^ is normally oxidized to Fe^3+^, which is bound to organic ligands in the environment (Hoffmann et al., 2012; Behnke and LaRoche, 2020). Understanding the availability of Fe in water is essential to understand how the Fe capture mechanism in the first algae may have developed initially.

**Figure 2.**
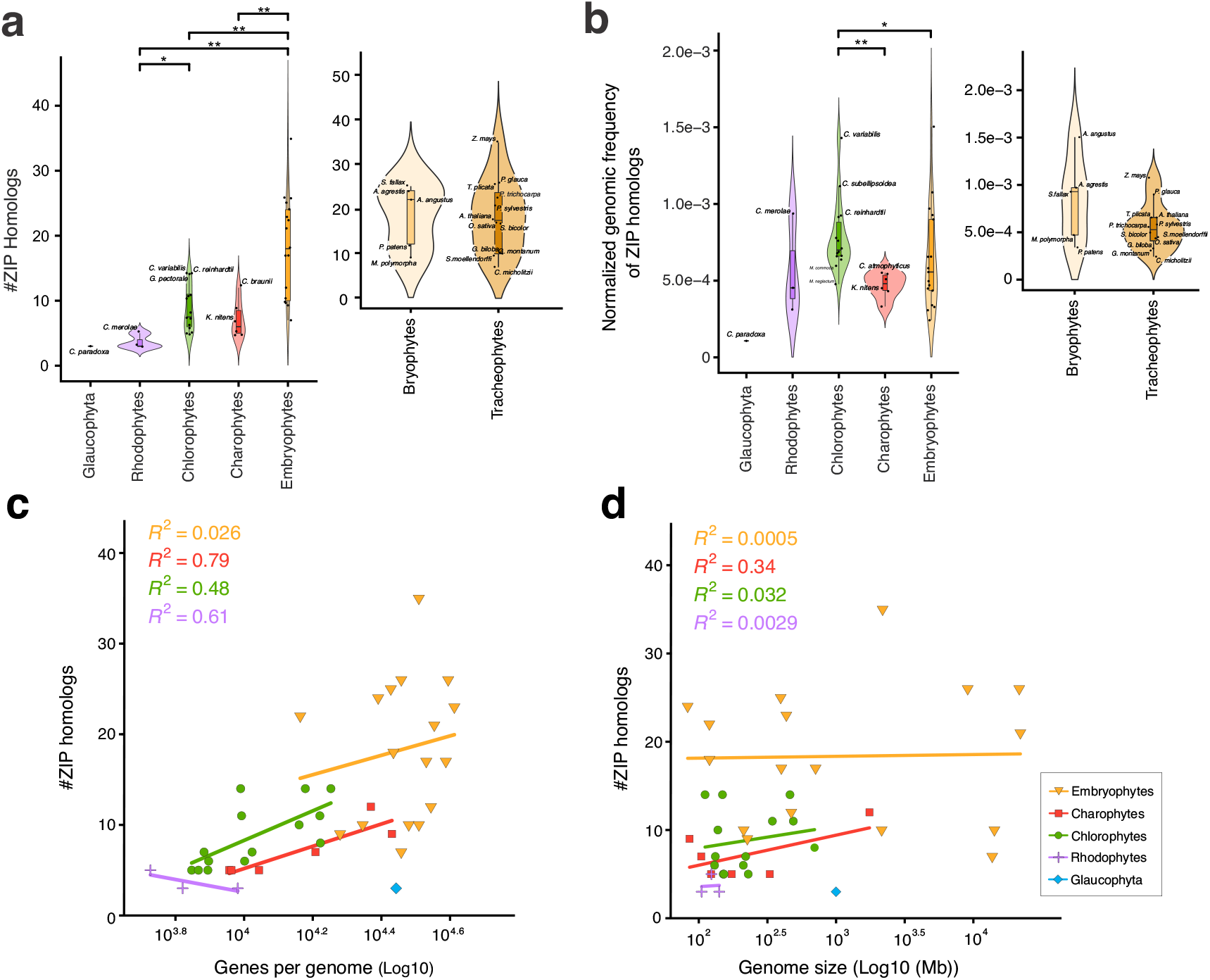
Distribution of *ZIP* genes across archaeplastidians. (a) Distribution of the number of ZIP homologs found per genome for diverse taxonomic groups. (b) Distribution of the relative genomic frequency of *ZIP* genes per genome (normalized by the total number of protein-coding genes per genome). ** represents p-value < 0.01 and * represents p-value < 0.05. Correlations between the number of *ZIP* genes per genome (c) and the total number of protein-coding genes per genome and (d) genome size.

Among archaeplastidians, land plants generally have a significantly greater number of *ZIP* genes (Figure 2a). However, we observed significant variability in the *ZIP* gene repertoire between land plant species, ranging from ~10 to ~50 genes per genome (Figure 2a). Little difference in the distribution of the number of *ZIP* homologs per genome was detected between non-vascular (Bryophyta) and vascular plants (Tracheophyta) (Figure 2a). It indicates that different lineages of bryophytes and tracheophytes might have experienced increased fitness provided by the fixation of additional copies of *ZIP* genes. A broader set of *ZIP* genes might be related to the evolution of more elaborated metal acquisition and internal transport due to compartmentalization of gene expression and metal specialization of the protein transporters.

The copy number of a gene family might be influenced by whole genome duplication (WGD), making it difficult to compare deeply divergent organisms like green plants, red algae and glaucophytes, that have experienced a variety of independent rounds of WGD. Thus, we decided to compare the relative genomic frequency of *ZIP* genes, calculated as the fraction of the total protein-coding gene set occupied by them. Results indicate that among green plants (Viridiplantae), chlorophytes tend to dedicate a significantly greater fraction of their gene set to encode ZIP proteins when compared to charophytes and land plants (Figure 2b). No significant difference in ZIP genomic frequency was detected regarding algal habitat (Supplemental Figure 2b), suggesting that no specific gene amplification trend can be linked to the habitat in archaeplastidians. Among chlorophytes, algae from the Class Chlorophyceae tend to have more *ZIP* genes per genome (Supplemental Figure 2c), although no differences regarding the relative genomic frequency of *ZIP* genes were found between the Chlorophyta Classes Chlorophyceae, Trebouxiophyceae, and Mamiellophyceae (Supplemental Figure 2d). In the unicellular freshwater chlorophytes *Chlorella variabilis* and *Coccomyxa subellipsoidea,* both from Class Trebouxiophyceae, > 0.1% of their protein-coding genes encode for ZIP proteins. A similar fraction is only observed in the top ZIP-enriched genomes of our land plant set such as maize and the hornwort *Anthoceros augustus* (Figure 2b). It is unclear why genomes of simple unicellular chlorophytes have accumulated such a high frequency of *ZIP* genes, more than double the median value observed for our set of vascular plants (~ 0.05%) (Figure 2b). *C. subellipsoidea* and *C. variabilis* are promising species for biofuel production and in the accumulation of metals such as cadmium (Cd) (Kováčik et al., 2017; Rana and Prajapati, 2021). In angiosperms it is known that some members of the ZIP family help in the absorption of Cd (Zheng et al., 2018). So, it is tempting to speculate that the large number of ZIP proteins in these algae might be related to the absorption of a broad range of metals other than iron.

No significant difference in genomic frequency of *ZIP* genes was detected between bryophytes and tracheophytes (Figure 2b), indicating that the variability of ZIP genomic frequency is similar between the two major groups of land plants. Although we accumulated knowledge in angiosperms such as *Arabidopsis* and rice, the molecular mechanisms of Fe acquisition are less well understood outside this group. Our results suggest, however, that *ZIP* genes might be important players in bryophytes. Experiments with the liverwort *Marchantia polymorpha* showed evidence for the use of Strategy I to capture Fe^2+^ through members of the ZIP family such as MpZIP3 (Mapoly0169s0010) and MpZIP5 (Mapoly0088s0041) (Lo et al., 2016).

We then wondered if the variation in the number of ZIP homologs between species could be correlated with other genomic features. We first compared the number of *ZIP* genes with the total number of protein-coding genes for each genome. Our results indicate that land plants do not exhibit any clear correlation between the two variables, suggesting that *ZIP* genes might not be preferably retained after WGD events in embryophytes (Figure 2c). The variation in the number of ZIP *loci* observed in land plants might be explained by gene-level duplications rather than large-scale gene duplication events. In opposition, data suggest that green algae species with more protein-coding genes tend to have more ZIP copies, indicating that independent expansions in the whole gene space of different taxa involved the fixation of duplicated copies of *ZIP* genes. Thus, the expanded set of ZIP observed in some green algae might be the result of gene fixation after lineage specific WGD events (Figure 2c). Unfortunately, the low number of available genomes of non-Viridiplantae algae prevents any further conclusions. Overall, *ZIP* genes occur in low copy number in the red algae and glaucophyte analyzed. Our data support the hypothesis that expansions in genome size *per se* might not explain the observed variation in *ZIP* genes between or within groups of archaeplastidians (Figure 2d).

### 3.2 Phylogenetic analysis of IRT1/ZIP proteins in Archaeplastida

To better understand the phylogenetic relationship between IRT1/ZIP proteins, we constructed a phylogenetic tree (Figure 3, Supplemental Figure 3) using the 492 unique *ZIP* genes identified across our sample of 41 genomes. The resulting phylogenetic tree can be subdivided into two deeply divergent major clades, called X and Y (Figure 3a). Interestingly, there was no clustering among the IRT1 proteins of land plants and green algae. IRT1/ZIP proteins from the angiosperms *Arabidopsis* and rice, and from the liverwort *Marchantia* all belong to Clade X, while *Chlamydomonas* IRT1 and IRT2 (herein called CrIRT1 and CrIRT2, respectively) belong to Clade Y (Figure 3a). MpZIP3 is phylogenetically close to the archetypical angiosperm IRT proteins from *Arabidopsis* and rice, while MpZIP5 belongs to a subclade more distantly related to *Arabidopsis* and rice IRTs that is shared with charophyte proteins. It indicates that divergent ZIP proteins might be recruited into Strategy I in land plants, possibly due to low specificity to metal transport and fast changes in regulatory networks that may include new *ZIP* genes in the early transcriptional response to environmental Fe depletion. We also performed a sequence similarity network (SSN) analysis with identical protein sequences that yielded similar results, confirming the distant relationship between Clades X and Y (Figure 3b). These results suggest that the mechanism of ferrous iron acquisition via IRT1/ZIP proteins evolved at least twice in green plants from deeply divergent ancestral genes of the ZIP family.

**Figure 3.**
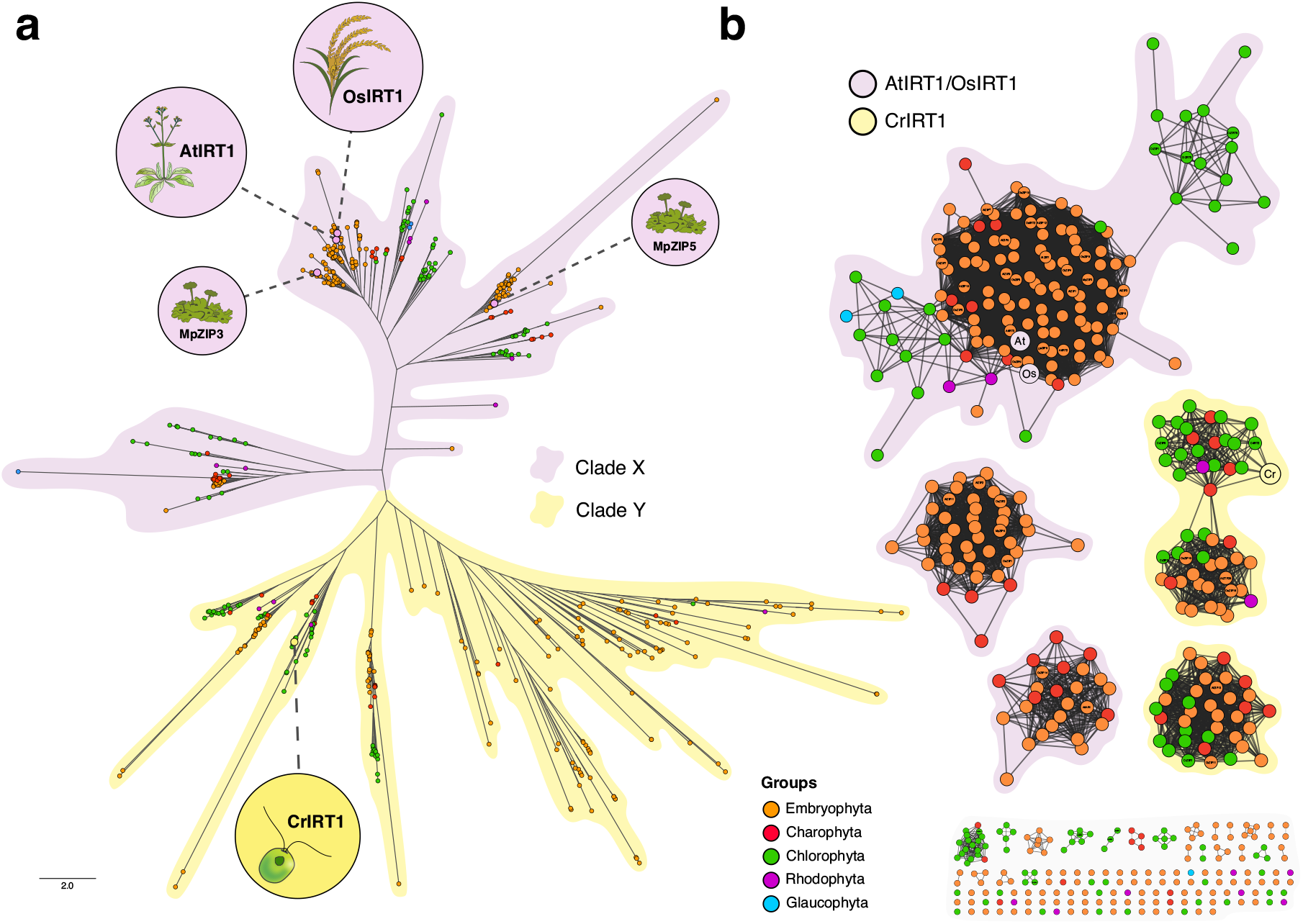
Maximum-likelihood and Sequence Similarity Network analyses of the IRT1/ZIP family in Archaeplastida. (a) Maximum-likelihood phylogenetic tree highlighting Clade X and Clade Y. Proteins discussed in the text are highlighted. (b) Sequence Similarity Networks showing clustering of proteins in agreement with the phylogenetic analysis. Color scheme reflects the taxonomic classification of species.

Angiosperm proteins AtZIP13 (AT3G08650), AtZTP29 (AT3G20870) and OsZIP13 (LOC_Os02g10230) from *Arabidopsis* and rice, respectively, are most likely Zn transporters that belong to Clade Y (Wang et al., 2010; Ivanov and Bauer, 2017; Zheng et al., 2018). It suggests that *Chlamydomonas* CrIRT1/2 are more closely related to Zn transporters proteins from angiosperms. The *Arabidopsis* proteins from Clade X AtZIP1 (AT3G12750) and AtZIP2 (AT5G59520) are involved in the uptake of Zn or Mn, or both (Milner et al., 2013; Zheng et al., 2018). Clade X proteins OsZIP5 (LOC_Os05g39560) and OsZIP9 (LOC_Os05g39540) function in Zn uptake from the soil in rice roots (Huang et al., 2020; Tan et al., 2020; Yang et al., 2020). The close related Clade X proteins AtIRT3 (AT1G60960), AtZIP4 (AT1G10970), and AtZIP9 (AT4G33020) from *Arabidopsis* and OsZIP7 (LOC_Os05g10940) from rice are involved in Zn xylem loading and root to shoot translocation (Ricachenevsky et al., 2018; Tan et al., 2019; Gindri et al., 2020; Lee et al., 2021). This suggests that the IRT1 from angiosperms like *Arabidopsis* and rice are more closely related to proteins acting in Zn or Mn transport than to the functional homologous CrIRT1 and CrIRT2 from *Chlamydomonas*.

These observations indicate that even deeply divergent ZIP proteins can evolve to fulfill the functional role of IRT1 in Fe^2+^ uptake first discovered in *Arabidopsis*. ZIP proteins might have a fast evolutionary turnover between different metal acquisition and transport mechanisms due to an overall low metal selectivity coupled with transcriptional network rewiring of *ZIP* genes (Nocedal and Johnson, 2015).

### 3.3 IRT1/ZIP protein variation

Previous works have demonstrated that the amino acid residues Aspartic Acid (D) at positions 108 and 144, Serine (S) at position 206 and Histidine (H) at position 232 of the *Arabidopsis* IRT1 are critical for iron transport (Figure 4a) (Rogers et al., 2000; Quintana et al., 2022). If those residues are substituted with Alanine (A), the protein loses its ability to transport Fe. These residues are also essential for the secondary transport of Zn in addition to Glutamic Acid (E) at position 111, which abolishes the transport of Zn when substituted with an A without affecting the transport of Fe (Figure 4a) (Rogers *et al.,* 2000). These results suggest that small changes in ZIP proteins can affect metal selectivity. However, if we align the CrIRT1 amino acid sequence with IRT1 from *Arabidopsis* and rice (Figure 4b) no conservation in three of the four critical positions is observed, suggesting that IRT proteins might exhibit some degree of sequence plasticity. Whether the presence of an A in the position 111 in the CrIRT1 is related to Zn transport activity should be better investigated with functional studies (Figure 4b).

**Figure 4.**
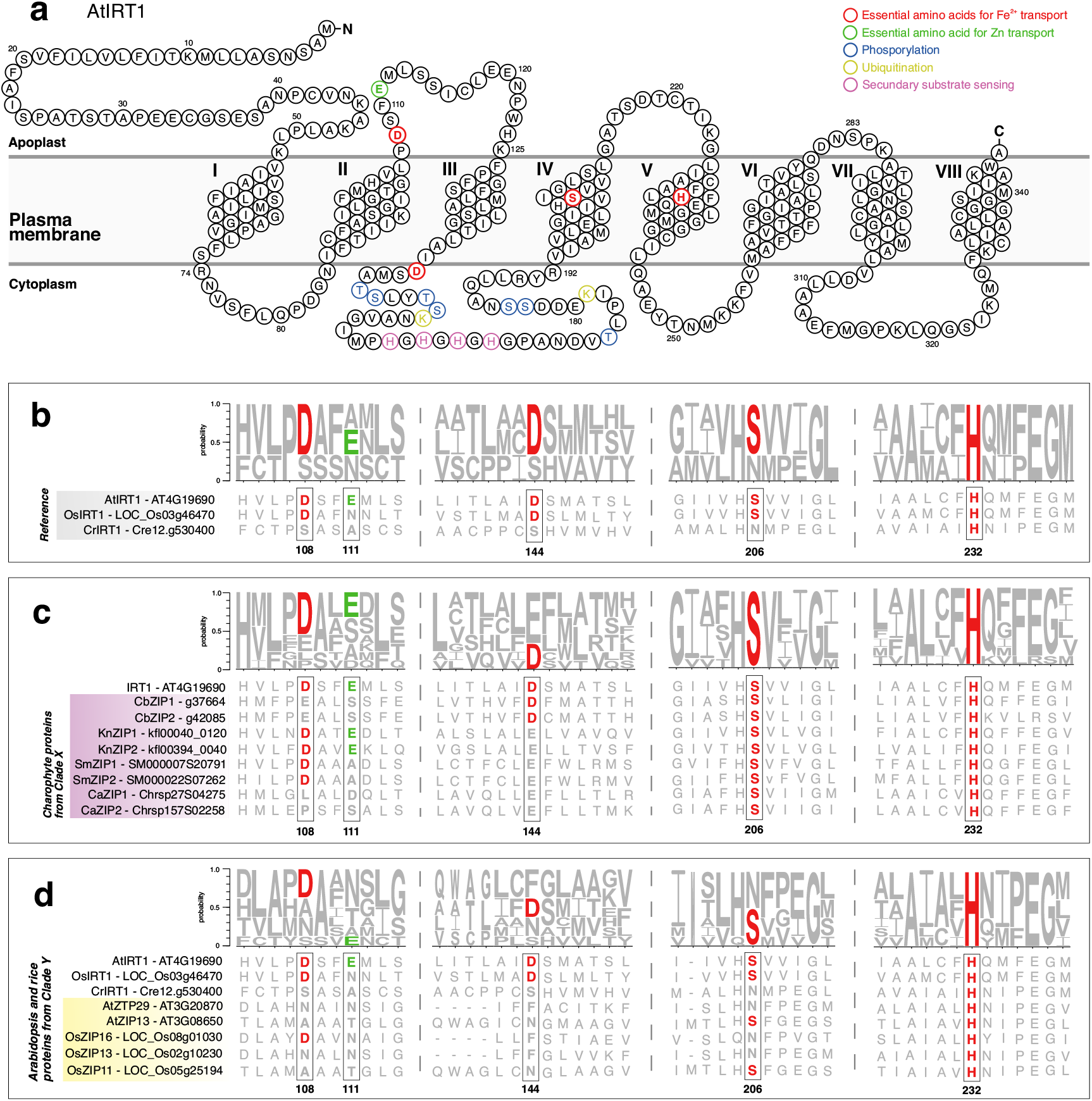
Multiple sequence alignments and logo sequence comparison of AtIRTl, OsIRTl and CrIRT1 with their closest related proteins. (a) Topology model of AtIRT1 highlighting the essential amino acid residues for Fe^2+^ transport (red), Zn transport (green), cytosolic phosphorylation (blue) and ubiquitination (yellow) sites, and H residues acting in secondary substrate sensing (purple). (b-d) Alignments showing the Asp-106, Asp-144, Ser-206 and His-232 (red) essential amino acids for Fe transport and Glu-111 (green) essential residue for Zn transport. The size of residues within a stack indicates the relative frequency of each amino acid per position. Protein amino acid numbering is related to AtIRT1. (b) Alignment between IRT1 sequences from Arabidopsis, rice and Chlamydomonas. (c) Charophyte ZIP proteins from Clade X aligned with the AtIRT1. (c) Arabidopsis and rice ZIP proteins from Clade Y aligned with AtIRT1, OsIRT1, and CrIRT1.

Our results suggest that Strategy I of land plants evolved based on Clade X IRT/ZIP proteins and could have evolved before the last common ancestor of Embryophyta. To better clarify this possibility, we aligned the *Arabidopsis* IRT1 with its closest related charophyte proteins from Clade X: g37664 (CbZIP1) and g42085 (CbZIP2) from *Chara braunii* (Charophyceae), kfl00040_0120 (KnZIP1) and kfl00394_0040 (KnZIP2) from *Klebsormidium nitens* (Klebsormidiophyceae), SM000007S20791 (SmZIP1) and SM000022S07262 (SmZIP2) from *Spirogloea muscicola* (Zygnematophyceae), and Chrsp27S04275 (CaZIP1) and Chrsp157S02258 (CaZIP2) from *Chlorokybus atmophyticus* (Chlorokybophyceae) (Figure 4c). We observed D108 position residue conservation in distinct charophyte lineages related to *Arabidopsis* IRT1. However, most charophyte IRT1-related proteins have an E residue in position 144 (Figure 4c). Interestingly, E and D residues are both acidic amino acids, with similar structure and chemical properties, indicating that the substitution D144E might not affect Fe transport. We also observed conservation in the essential amino acids S206 and H232, which are reported to be critical for its *in planta* function of high-affinity Fe^2+^ transport activity (Quintana et al., 2022), in all analyzed charophyte proteins (Figure 4c). These results suggest that the charophyte proteins from Clade X closely related to IRT1 from *Arabidopsis* and rice are compatible with Fe^2+^ acquisition in Strategy I and its role in Fe homeostasis. It opens the possibility that Strategy I might have evolved during early land colonization by simple charophytes before the origin of land plants. Functional studies in charophytes, especially with terrestrial/subaerial species, will be needed to confirm this hypothesis.

To check whether Clade Y proteins have evolved diverse metal affinities, we aligned the *Chlamydomonas* IRT1 with its closest related proteins from *Arabidopsis* and rice (AtZTP29, AtZIP13, OsZIP13, OsZIP16, and OsZIP11), together with IRT1 from *Arabidopsis* and rice from Clade X (Figure 4d). No apparent sequence conservation was observed in the critical residues, except for H232, suggesting that Clade Y and Clade X proteins might have relevant amino acid sequence differences and still be able to function in the transport of the same metal ions.

We next wondered about structural variations in ZIP proteins, related to the presence of additional protein domains. Our results indicate that ~90% of the ZIP proteins of archaeplastidians present only the ZIP domain (PF02535) as seen for the archetypical IRT1 proteins from *Arabidopsis* and rice. The remaining ~10% present the ZIP domain associated with at least another domain. In rare cases, the ZIP domain was associated with tens of different domains in the same predicted protein (Supplemental Figure 4a). Overall, land plants and chlorophytes tend to have ZIP proteins associated with other domains more frequently than other taxa (Supplemental Figure 4b). The domain Presenilin (PF01080) is found associated with the ZIP domain in 34 proteins (~6.9% of the sample), while the domains NPR3 (PF03666) and CDC45-like (PF02724) in 25 (~5.1%) and 24 proteins (~4.9%), respectively (Supplemental Figure 4c). Some ZIP proteins present combinations between those domains (Supplemental Figure 4d). The Presenilin domain is usually associated with transmembrane proteins (Hass et al., 2009; Mistry et al., 2021) and NPR3 domain was described as a regulator of Target of Rapamycin-TOR Complex 1 (TORC1), a central regulator of eukaryotes cell growth, in response to amino acid starvation (Neklesa and Davis, 2009; Wu and Tu, 2011). CDC45 is involved in the initiation of DNA replication in *S. cerevisiae* (Saha *et al.,* 1998), although not much is known about CDC45-like proteins in plants. The role of the ZIP domain in such multidomain proteins remains to be further investigated.

### 3.4 Structural analyses of IRT1/ZIP proteins

To further support the hypothesis that divergent ZIP proteins fulfill the role of Fe^2+^ performed by IRT1, we modeled the tridimensional structure of the Fe uptaking protein from land plants (AtIRT1, OsIRT1, MpZIP3 and MpZIP5 from Clade X) and the chlorophyte *Chlamydomonas* (CrIRT1 from Clade Y). We also included in the analysis the Clade X charophyte proteins CbZIP1, KnZIP1, SmZIP1, CaZIP1 and the Clade Y putative Zn transporters OsZIP13 and AtZTP29 (Supplemental Table 2). The predicted structures were aligned in pairs, allowing us to estimate the structural similarity between proteins.

The pairwise RMSD values suggest that, Clade X proteins are structurally conserved (Figure 5a). The angiosperm proteins AtIRT1 and OsIRT1 have highly similar predicted structures, with an RMSD value of 0.53 Å (lower values indicate higher structural similarity). Interestingly, the charophyte proteins from Clade X have a high degree of structural similarity with AtIRT1, with pairwise RMSD values ranging between 0.79 and 1.68 Å (Figure 5a and 5b), suggesting that they might be functionally conserved. The IRT/ZIP proteins from the soil charophytes *Klebsormidium* (KnZIP1) and *Spirogloea* (SmZIP1) showed remarkable structural conservation with AtIRT1 (RMSD = 0.79 and 1.08 Å, respectively), suggesting that Strategy I Fe uptaking via IRT1 might have evolved in the Streptophyta lineage before the origin of land plants, during the adaptation of early charophytes to the Fe acquisition in terrestrial settings.

**Figure 5.**
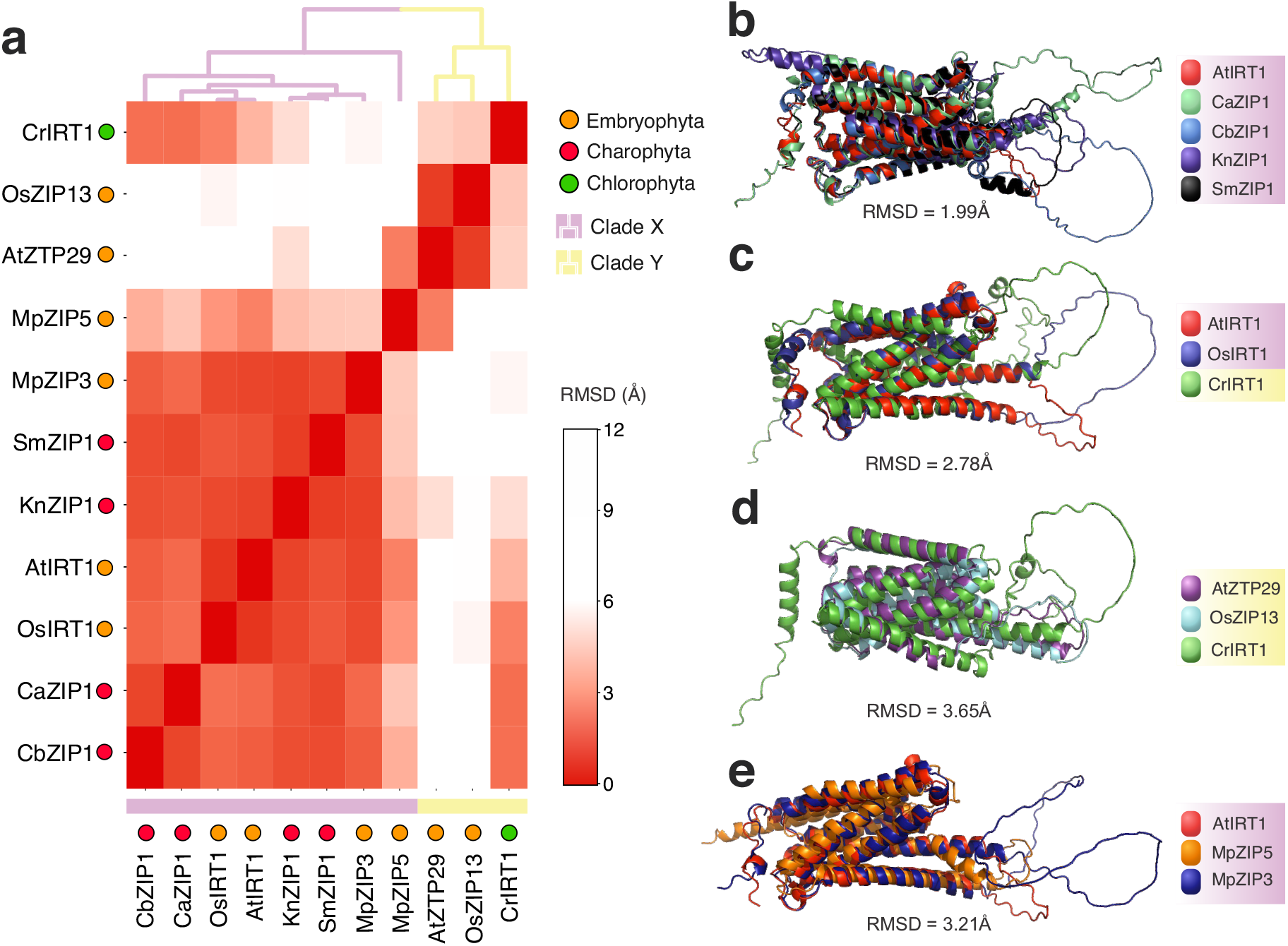
IRT1/ZIP tridimensional structure analysis. (a) Heatmap showing the structural similarity of pairwise alignments between selected IRT1/ZIP proteins from Clade X and Clade Y. The structural similarity is indicated by the RMSD (Å) value, where the lower value (red) indicates more structural similarity. (b-e) Aligned structural models of (b) CrIRT1, AtIRT1 and OsIRT1; (c) AtIRT1 and its closest related charophyte ZIP proteins from clade X; (d) CrIRT1, AtZTP29 and OsZIP13 from clade Y; and (e) MpZIP5, MpZIP3 and AtIRT1.

Surprisingly, the predicted structure of the Fe transporter CrIRT1 from the chlorophyte *Chlamydomonas* (Clade Y) showed more similarity to structures from the Clade X, such as the Fe transporters OsIRT1 (RMSD = 2.2 Å) and AtIRT1 (RMSD = 3.8 Å) (Figure 5a and 5c), than to the phylogenetically related putative Zn transporters OsZIP13 (RMSD = 4.3 Å) and AtZTP29 (RMSD = 4.6 Å) from Clade Y (Figure 5a and 5d). These results indicate that, although CrIRT1 and AtIRT1/OsIRT1 are only distantly related in terms of ZIP family phylogeny, as revealed by their primary sequence of amino acids (Figure 3), they evolved to some degree of structure convergence which might be related to the Fe transport activity. While the predicted structure of the Fe transporter MpZIP3 from the liverwort *Marchantia* is highly similar to other ZIP proteins from the Clade X (RMSD ranging from 0.7 to 1.6 Å), MpZIP5 structure shows a more moderate structural conservation with proteins from both clades Y (RMSD with AtZTP29 = 2.2 Å) and X (RMSD with AtIRT1 = 2.3 Å) (Figure 5a and 5e). These results indicate that ZIP proteins involved in the acquisition of Fe^2+^ can tolerate some degree of structural variations and still be functional.

All Clade X and Y protein models showed a cytosolic loop of variable length (Figure 5b, 5c, 5d & 5e) that is consistently found between transmembrane helices III and IV, as seen for AtIRT1(Figure 4a; Supplemental Table 4). CrIRT1 is the only protein analyzed where the cytosolic loop is found between transmembrane helices V and VI (Supplemental Table 4). Specific amino acids in this loop are target of post -translational protein modifications in *Arabidopsis* (Figure 4a), being involved in protein ubiquitination, internalization, and degradation, as part of the mechanism of control of Fe homeostasis (Barberon et al., 2011; Quintana et al., 2022). We were unable to find apparent conservation of such amino acids between the modeled IRT1/ZIP proteins, indicating that the loop sequence might be variable (Supplemental Table 4). In fact, the loop sequences are significantly more variable than the regular protein structural motifs (Supplemental Figure 5), suggesting that this regulatory portion of the protein might evolve fast between IRT1/ZIP homologs. Our results indicate that post-translational protein modifications controlling IRT1/ZIP protein turnover are likely variable between taxa.

## 4. Conclusions

Our results support the notion that green plants functional homologs of the archetypical angiosperm IRT1 emerged from distantly related ZIP proteins that are divergent in key features of their amino acid sequences. Furthermore, the typical critical amino acids in the *Arabidopsis* IRT1 protein seems to have evolved before the split of land plants from streptophytic algae. In fact, structural modeling suggests that putative IRT1 proteins from charophytes are highly conserved with the archetypical angiosperm IRT1 counterparts, indicating that the IRT1-dependent Fe uptake typically found among land plants might have originated in early terrestrial charophytes before the origin of the first embryophytes. Functional data on Fe acquisition of soil charophytes (such as *Klebsormidium, Spirogloea* and *Mesotaenium)* might help understand how land plants ancestors were able to first obtain Fe in terrestrial settings with subaerial exposure and direct contact with the substrate. On the other hand, IRT1/2 from the chlorophyte *Chlamydomonas* has a divergent protein sequence regarding key residues known in *Arabidopsis*, suggesting that multiple primary amino acid sequences of ZIP proteins might be fitted to transport Fe^2+^ efficiently. This might be explained, at least partially, by the evolution of a structural convergence between distantly related IRT1/ZIP proteins.

The phylogenetic diversity of IRT1/ZIP proteins used in Fe uptake by green plants indicate that networking rewiring of *ZIP* genes encoding proteins with the ability to transport Fe might be the ultimate cause of IRT1 diversity in the Strategy I of Fe acquisition. Besides its ecological importance for photosynthesis, little is known about Fe acquisition strategies across the superdiverse group Archaeplastida. The results presented here indicate that functional homologs of IRT1 can emerge from highly divergent phylogenetic groups of ZIP metal transporters, suggesting that much evolutionary diversity is still to be found in Fe uptake in less studied groups such as green and red algae.

## 5. Acknowledgments

This work was supported by the Serrapilheira Institute (grant number Serra-1812-26691) and by the Coordenação de Aperfeiçoamento de Pessoal de Nível Superior - Brasil (CAPES) – Finance Code 001, that provided a master’s and PhD scholarship to WFCR.

## 6. Author Contributions

LEDB designed the research. WFCR carried out the sequence data collection, analyses and visualizations. ABPL contributed to protein three-dimensional modeling and structural analyses. LEDB, WFCR, FKR and JEL contributed to data interpretation and manuscript structure. WFCR and LEDB wrote the manuscript. All authors participated in manuscript editing and approved the final version.

## Supporting Information Legends

**Supplemental Figure 1. Phylogeny of Archaeplastida.** Faded colors (orange, red, green, purple, and blue) represent the main archaeplastidian groups analyzed in this work. Topology was based on Lemieux et al., (2015), Cole et al., (2019a,b), Li et al., (2020), Wang et al., (2021).

**Supplemental Figure 2. Distribution of ZIP genes across algae.** Plots showing the global distribution of ZIP homologs, and the frequency of proteins found normalized by the number of genes per genome by algae species in different habitats (a, b) and different Chlorophyta class (c, d).

**Supplemental Figure 3. Maximum-likelihood phylogenetic tree of ZIP family in Archaeplastida.** Complete circular phylogenetic tree with full protein IDs containing all 492 ZIP proteins found in our dataset with 41 archaeplastidian species.

**Supplemental Figure 4. Domain analysis of ZIP proteins found in Archaeplastida.** (a) Network summarizing the sequences (squares) connected to their respective protein domains (circles). In the network, the domains that have the most connections with protein sequences stand out. (b) Distribution of the number of unique domains per sequence in Archaeplastida groups and their medians. Sequences with more than 6 domains were discarded. (c) Bar plot showing the 20 most common domains that are found associated with the ZIP domain by the number of occurrences in different proteins. (d) Venn diagram showing the occurrence of multidomain proteins and the dominance of single ZIP domain in ~90% of ZIP proteins.

**Supplemental Figure 5. Pairwise distances of the modeled proteins.** Boxplots showing the sequence distances calculated by p-distance (number of differences/number of aligned positions) between protein cytosolic loops and the rest of the protein sequences. Briefly, p-distance = 0.0 indicates total conservation and p-distance =1.0 indicates no sequence conservation. Analysis indicate that the loop portion of the IRT1/ZIP proteins is more variable than the rest of the proteins. The black dots indicate the means and red bars are the standard deviation.

**Supplemental Table 1.** List of complete predicted proteome databases used in this study.

**Supplemental Table 2.** IRT1/ZIP Proteins used in the structural modelling. If predicted, signal peptides were removed from the final protein model.

**Supplemental Table 3.** Table showing the number of ZIP family homologs found by species in our dataset.

**Supplemental Table 4.** Cytosolic loops identified in the modeled IRT1/ZIP proteins.

