## Supplementary figures and images for "Ferrous iron uptake via IRT1/ZIP evolved at least twice in green plants"

### Supplemental Figure 1

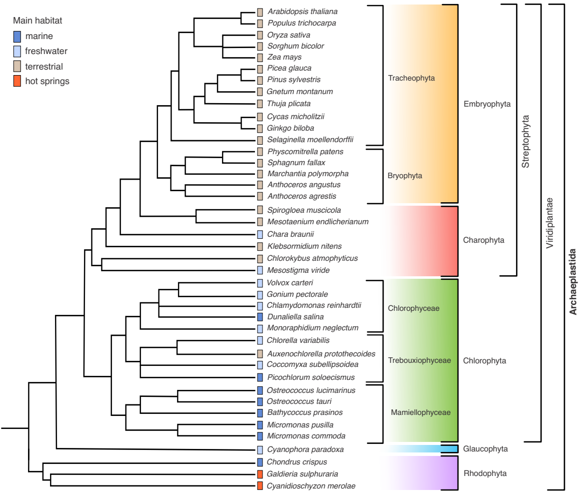

### Supplemental Figure 2

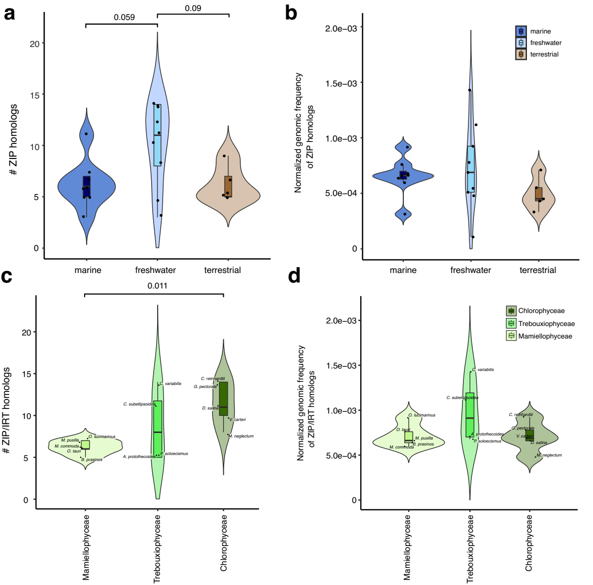

### Supplemental Figure 3

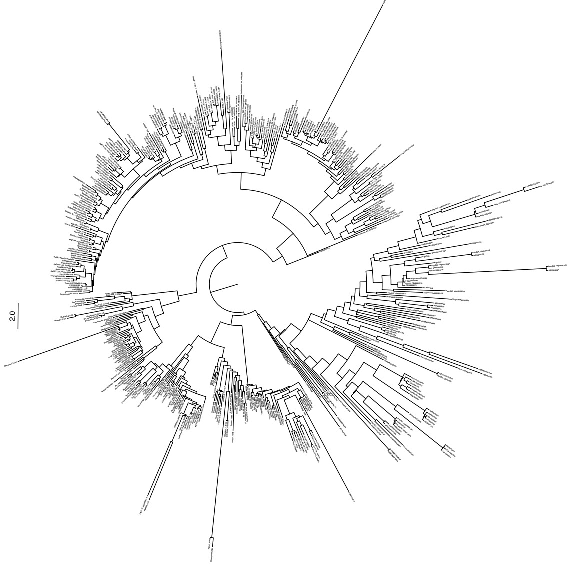

### Supplemental Figure 4

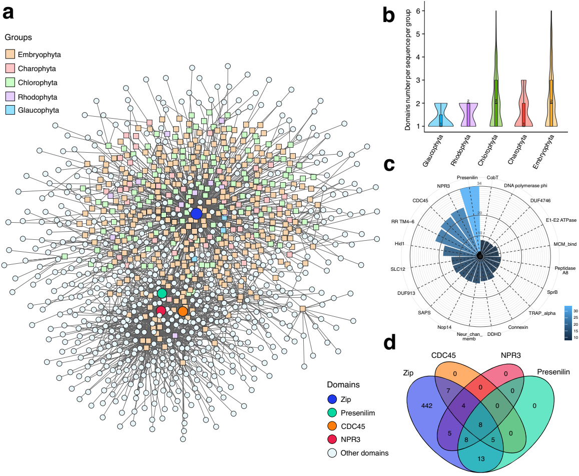

### Supplemental Figure 5

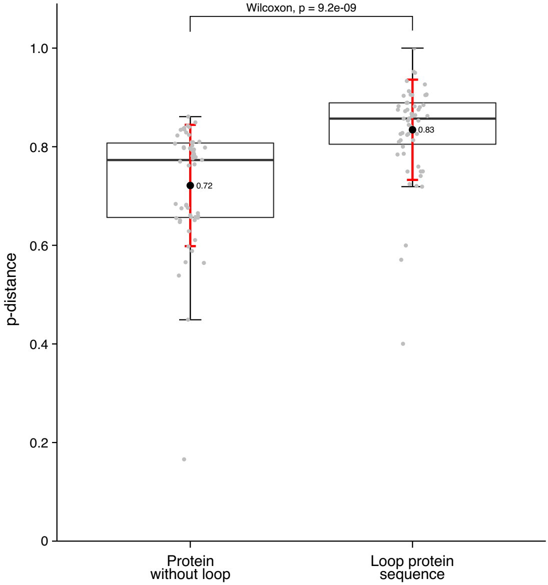
