## Supplemental Table 1 for "Ferrous iron uptake via IRT1/ZIP evolved at least twice in green plants"

**Supplemental Table S1**. List of complete predicted proteome databases used in this study.

| **Species** | **Group** | **Link** |
| --- | --- | --- |
| *Arabidopsis thaliana* | Tracheophyta | https://phytozome-next.jgi.doe.gov/info/Athaliana_TAIR10 |
| *Oryza sativa* | Tracheophyta | https://phytozome-next.jgi.doe.gov/info/Osativa_v7_0 |
| *Populus trichocarpa* | Tracheophyta | https://phytozome-next.jgi.doe.gov/info/Ptrichocarpa_v4_1 |
| *Sorghum bicolor* | Tracheophyta | https://phytozome-next.jgi.doe.gov/info/Sbicolor_v3_1_1 |
| *Selaginella moellendorffii* | Tracheophyta | https://phytozome-next.jgi.doe.gov/info/Smoellendorffii_v1_0 |
| *Zea mays* | Tracheophyta | https://phytozome-next.jgi.doe.gov/info/Zmays_RefGen_V4 |
| *Ginkgo biloba* | Tracheophyta | ftp://ftp.psb.ugent.be/pub/plaza/plaza_gymno_01/Fasta/proteome.gbi.csv.gz |
| *Cycas micholitzii* | Tracheophyta | ftp://ftp.psb.ugent.be/pub/plaza/plaza_gymno_01/Fasta/proteome.cmi.csv.gz |
| *Picea glauca* | Tracheophyta | ftp://ftp.psb.ugent.be/pub/plaza/plaza_gymno_01/Fasta/proteome.pgl.csv.gz |
| *Pinus sylvestris* | Tracheophyta | ftp://ftp.psb.ugent.be/pub/plaza/plaza_gymno_01/Fasta/proteome.psy.csv.gz |
| *Thuja plicata* | Tracheophyta | ttps://phytozome-next.jgi.doe.gov/info/Tplicata_v3_1 |
| *Gnetum montanum* | Tracheophyta | ftp://ftp.psb.ugent.be/pub/plaza/plaza_gymno_01/Fasta/proteome.gmo.csv.gz |
| *Anthoceros agrestis [Oxford]* | Bryophyta | https://www.hornworts.uzh.ch/static/download/a_agr_oxford.zip |
| *Anthoceros angustus* | Bryophyta | https://datadryad.org/stash/dataset/doi:10.5061/dryad.msbcc2ftv |
| *Marchantia polymorpha* | Bryophyta | https://phytozome-next.jgi.doe.gov/info/Mpolymorpha_v3_1 |
| *Sphagnum fallax* | Bryophyta | https://phytozome-next.jgi.doe.gov/info/Sfallax_v1_1 |
| *Physcomitrella patens* | Bryophyta | https://phytozome-next.jgi.doe.gov/info/Ppatens_v3_3 |
| *Chara braunii* | Charophyta | https://bioinformatics.psb.ugent.be/gdb/Chara_braunii/chbra_iso_noTE_23546_pep.fasta |
| *Klebsormidium nitens* | Charophyta | http://www.plantmorphogenesis.bio.titech.ac.jp/~algae_genome_project/klebsormidium/kf_download/160614_klebsormidium_v1.1_AA.fasta |
| *Mesotaenium endlicherianum* | Charophyta | https://ndownloader.figshare.com/files/17819138 |
| *Chlorokybus atmophyticus* | Charophyta | ftp://ftp.cngb.org/pub/CNSA/data1/CNP0000228/CNS0021447/CNA0002353/scaffold_Chlorokybus_atmophyticus.pep.gz |
| *Mesostigma viride* | Charophyta | ftp://ftp.cngb.org/pub/CNSA/data1/CNP0000228/CNS0021438/CNA0002352/scaffold_Mesostigma_viride.pep.gz |
| *Spirogloea muscicola* | Charophyta | https://ndownloader.figshare.com/files/17819147 |
| *Auxenochlorella protothecoides* | Chlorophyta | https://genome.jgi.doe.gov/portal/Auxeprot1/download/Auxeprot1_GeneCatalog_proteins_20170909.aa.fasta.gz |
| *Bathycoccus prasinos* | Chlorophyta | https://genome.jgi.doe.gov/portal/Batpra1/download/Batpra1_GeneCatalog_proteins_20180426.aa.fasta.gz |
| *Chlorella variabilis* | Chlorophyta | https://genome.jgi.doe.gov/portal/ChlNC64A_1/download/Chlorella_NC64A.all_proteins.fasta.gz |
| *Coccomyxa subellipsoidea* | Chlorophyta | https://genome.jgi.doe.gov/portal/Coc_C169_1/download/Coccomyxa_C169_v2_all_proteins.fasta.gz |
| *Chlamydomonas reinhardtii* | Chlorophyta | https://genome.jgi.doe.gov/portal/pages/dynamicOrganismDownload.jsf?organism=Phytozome# |
| *Dunaliella salina* | Chlorophyta | https://genome.jgi.doe.gov/portal/Dunsal1/download/Dunsal1_GeneCatalog_proteins_20180508.aa.fasta.gz |
| *Gonium pectorale* | Chlorophyta | https://genome.jgi.doe.gov/portal/Gonpec1/download/Gonpec1_GeneCatalog_proteins_20180501.aa.fasta.gz |
| *Micromonas commoda* | Chlorophyta | https://genome.jgi.doe.gov/portal/MicpuN3v2/download/MicpuN3v2_GeneCatalog_proteins_20160404.aa.fasta.gz |
| *Monoraphidium neglectum* | Chlorophyta | https://genome.jgi.doe.gov/portal/Monneg1/download/Monneg1_GeneCatalog_proteins_20170920.aa.fasta.gz |
| *Micromonas pusilla* | Chlorophyta | https://genome.jgi.doe.gov/portal/MicpuC2/download/MicromonasCCMP1545.allModels.aa.fasta.gz |
| *Ostreococcus lucimarinus* | Chlorophyta | https://phytozome-next.jgi.doe.gov/info/Olucimarinus_v2_0 |
| *Ostreococcus tauri* | Chlorophyta | https://genome.jgi.doe.gov/portal/Ostta4221_3/download/Ostta4221_3_GeneCatalog_proteins_20161028.aa.fasta.gz |
| *Picochlorum soloecismus* | Chlorophyta | https://genome.jgi.doe.gov/portal/Picsp_1/download/Picsp_1_GeneCatalog_proteins_20170909.aa.fasta.gz |
| *Volvox carteri* | Chlorophyta | https://phytozome-next.jgi.doe.gov/info/Vcarteri_v2_1 |
| *Chondrus crispus* | Rhodophyta | https://greenhouse.lanl.gov/greenhouse/organisms/ |
| *Cyanidioschyzon merolae* | Rhodophyta | https://greenhouse.lanl.gov/greenhouse/organisms/ |
| *Galdieria sulphuraria* | Rhodophyta | https://greenhouse.lanl.gov/greenhouse/organisms/ |
| *Cyanophora paradoxa* | Glaucophyta | https://greenhouse.lanl.gov/greenhouse/organisms/ |
