## Supplemental Table 2 for "Ferrous iron uptake via IRT1/ZIP evolved at least twice in green plants"

**Supplemental Table S2.** IRT1/ZIP Proteins used in the structural modeling. If predicted, signal peptides were removed from the final protein model.

| **Protein name** | **Organism** | **Protein ID** | **Accession code** | **Signal peptide (aa position)** | **Protein length (aa)** |
| --- | --- | --- | --- | --- | --- |
| AtIRT1 | *Arabidopsis thaliana* | AT4G19690 | Q38856**^A^** | 1-27 | 347 |
| CrIRT1 | *Chlamydomonas reinhardtii* | Cre12.g530400 | A0A2K3D4S5**^A^** | 1-28 | 367 |
| OsIRT1 | *Oryza sativa* | LOC_Os03g46470 | Q75HB1**^A^** | 1-33 | 374 |
| CbZIP1 | *Chara braunii* | g37664 | A0A388LNG6**^A^** | NA | 375 |
| KnZIP1 | *Klebsormidium nitens* | kfl00040_0120 | GAQ79869.1**^B^** | NA | 340 |
| SmZIP1 | *Spirogloea muscicola* | SM000007S20791 | NA | NA | 319 |
| CaZIP1 | *Chlorokybus atmophyticus* | Chrsp27S04275 | NA | NA | 381 |
| AtZTP29 | *Arabidopsis thaliana* | AT3G20870 | Q940Q3**^A^** | 1-22 | 276 |
| OsZIP13 | *Oryza sativa* | LOC_Os02g10230 | EEE56500.1**^B^** | NA | 276 |
| MpZIP3 | *Marchantia polymorpha* | Mapoly0169s0010 | A0A2R6W308**^A^** | NA | 381 |
| MpZIP5 | *Marchantia polymorpha* | Mapoly0088s0041 | A0A0E3DAH5**^A^** | 1-32 | 344 |

**^A^:** UNIPROT accession code.

**^B^**: GENBANK accession code.
