## Supplemental Table 3 for "Ferrous iron uptake via IRT1/ZIP evolved at least twice in green plants"

**Supplemental Table S3**. Table showing the number of ZIP family homologs found by species in our dataset.

| Species | Group | # of ZIP homologs |
| --- | --- | --- |
| *Arabidopsis thaliana* | Embryophyta | 18 |
| *Marchantia polymorpha* | Embryophyta | 9 |
| *Oryza sativa* | Embryophyta | 17 |
| *Zea mays* | Embryophyta | 35 |
| *Physcomitrella patens* | Embryophyta | 12 |
| *Populus trichocarpa* | Embryophyta | 23 |
| *Sorghum bicolor* | Embryophyta | 17 |
| *Selaginella moellendorffii* | Embryophyta | 10 |
| *Anthoceros agrestis* [Oxford] | Embryophyta | 24 |
| *Sphagnum fallax* | Embryophyta | 25 |
| *Anthoceros angustus* | Embryophyta | 22 |
| *Ginkgo biloba* | Embryophyta | 10 |
| *Cycas micholitzii* | Embryophyta | 7 |
| *Picea glauca* | Embryophyta | 26 |
| *Pinus sylvestris* | Embryophyta | 21 |
| *Thuja plicata* | Embryophyta | 26 |
| *Gnetum montanum* | Embryophyta | 10 |
| *Chlorokybus atmophyticus* | Charophyta | 5 |
| *Chara braunii* | Charophyta | 12 |
| *Klebsormidium nitens* | Charophyta | 7 |
| *Mesotaenium endlicherianum* | Charophyta | 5 |
| *Mesostigma viride* | Charophyta | 5 |
| *Spirogloea muscicola* | Charophyta | 9 |
| *Auxenochlorella protothecoides* | Chlorophyta | 5 |
| *Bathycoccus prasinos* | Chlorophyta | 5 |
| *Chlorella variabilis* | Chlorophyta | 14 |
| *Coccomyxa subellipsoidea* | Chlorophyta | 11 |
| *Chlamydomonas reinhardtii* | Chlorophyta | 14 |
| *Dunaliella salina* | Chlorophyta | 11 |
| *Gonium pectorale* | Chlorophyta | 14 |
| *Micromonas commoda* | Chlorophyta | 6 |
| *Monoraphidium neglectum* | Chlorophyta | 8 |
| *Micromonas pusilla* | Chlorophyta | 7 |
| *Ostreococcus lucimarinus* | Chlorophyta | 7 |
| *Ostreococcus tauri* | Chlorophyta | 6 |
| *Picochlorum soloecismus* | Chlorophyta | 5 |
| *Volvox carteri* | Chlorophyta | 10 |
| *Chondrus crispus* | Rhodophyta | 3 |
| *Cyanidioschyzon merolae* | Rhodophyta | 5 |
| *Galdieria sulphuraria* | Rhodophyta | 3 |
| *Cyanophora paradoxa* | Glaucophyta | 3 |
| Total |  | 492 |
