## Supplemental Table 4 for "Ferrous iron uptake via IRT1/ZIP evolved at least twice in green plants"

**Supplemental Table S4**. Cytosolic loops identified in the modeled IRT1/ZIP proteins.

| **Protein name** | **Loop length** | **Loop between helixes** | **Loop sequence** |
| --- | --- | --- | --- |
| AtIRT1 | 158-180 | 3-4 | GIMPHGHGHGHGPANDVTLPIKE |
| CrIRT1 | 87-166 | 5-6 | HGSMLASGSSFCTPSASASCSSCCCTGGEAHID  EKSAAACPPCSHVMVHVAPQPVASGDGEDVMAGGGGK  QAGGRAGSSA |
| OsIRT1 | 163-205 | 3-4 | RSKPRPSSGGDVAAVADHGESPDQGHRHGHGHGHGHGMAVAKP |
| CbZIP1 | 179-205 | 3-4 | GAGAMLAVTAMGKTPGAEGKGDVVVDC |
| KaZIP1 | 130-182 | 3-4 | GHHHPDHVHVHGLAHQGVAPSVDSGDAHPPAGSVKSLDVEAG  SVAKESTFGTP |
| SmZIP1 | 119-150 | 3-4 | YGLPTQASLEDSDEVQGGLSMGDDLEGAKANP |
| CaZIP1 | 164-200 | 3-4 | LAQDVHQPTVPSTSTTTTITSSAAPTLVHRGHAHGAG |
| AtZTP29 | 96-106 | 2-3 | FIPEPTLGPSTDGKRRKKNGD |
| OsZIP13 | 86-108 | 3-4 | FIPEPTVVPTADAGKKQTDDDGS |
| MpZIP3 | 130-209 | 3-4 | SHGHTEDYEKLPTRKSILDSDSTHEPLLQNSSTSLAVAEPVHKTN  GSQHHEKHTERSHTHTPNGGADEEAGHNGNHLVLA |
| MpZIP5 | 157-188 | 3-4 | TGKESAITGAVGPNDHEAGHMVQATLIHTASM |
